# Absence of SARS-CoV-2 infection in cats and dogs in close contact with a cluster of COVID-19 patients in a veterinary campus

**DOI:** 10.1101/2020.04.07.029090

**Authors:** Sarah Temmam, Alix Barbarino, Djérène Maso, Sylvie Behillil, Vincent Enouf, Christèle Huon, Ambre Jaraud, Lucie Chevallier, Marija Backovic, Philippe Pérot, Patrick Verwaerde, Laurent Tiret, Sylvie van der Werf, Marc Eloit

## Abstract

Severe acute respiratory syndrome coronavirus 2 (SARS-CoV-2), which originated in Wuhan, China, in 2019, is responsible for the COVID-19 pandemic. It is now accepted that the wild fauna, probably bats, constitute the initial reservoir of the virus, but little is known about the role pets can play in the spread of the disease in human communities, knowing the ability of SARS-CoV-2 to infect some domestic animals. We tested 21 domestic pets (9 cats and 12 dogs) living in close contact with their owners (belonging to a veterinary community of 20 students) in which two students tested positive for COVID-19 and several others (n = 11/18) consecutively showed clinical signs (fever, cough, anosmia, etc.) compatible with COVID-19 infection. Although a few pets presented many clinical signs indicative for a coronavirus infection, no animal tested positive for SARS-CoV-2 by RT-PCR and no antibodies against SARS-CoV-2 were detectable in their blood using an immunoprecipitation assay. These original data can serve a better evaluation of the host range of SARS-CoV-2 in natural environment exposure conditions.

---

SARS-CoV-2 infection has presented unprecedented challenges related to viral disease control and prevention worldwide. While the emergence of the virus is now well-documented, important questions regarding the transmissibility of the disease among human populations remain to be answered. In particular, the hypothesis that small domestic animals could serve as intermediate or amplification hosts of the virus needs to be further addressed, in order to improve epidemiological models and to undertake appropriate preventive measures.

We investigated the presence of SARS-CoV-2 infection of domestic cats (n = 9) and dogs (n = 12) living in close contact with French veterinary students who were their owners (n = 18), belonging to a community in which 11 of them (11/18; 61%) developed symptoms compatible with COVID-19 between February 25^th^ and March 18^th^, 2020, including two students were tested positive for SARS-CoV-2 by RT-PCR. The other 9 patients were not tested, in accordance with current French regulations, which do not prescribe to test all patients in such clusters. Other owners (7/18; 39%) were asymptomatic at the time of sampling. All of the owners were in the close vicinity of their pets, *e*.*g*. living in the same room – of 12 to 17 square meters - and sharing the same bed (100% of cats’ owners and 33% of dog’s owners); accepting face/hand licking (78% and 92% of cats’ and dogs’ owners, respectively). All cats were domestic European Shorthair cats. Dogs included in the sampled panel were either cross-bred (n = 6) or purebred individuals from the Labrador Retriever, Shetland Sheepdog, Belgian Malinois and White Swiss Shepherd breeds (n = 6). The average age of these sampled adult animals was 3.3 years for cats (min: 6 months; max: 6.5 years) and 2.7 years for dogs (min: 4 months; max: 8 years). Small pets were free of clinical signs, except some respiratory or digestive signs reported for 3 cats, independently.

Sera from this per-epidemic cohort of pets (hereafter named “pets epidemic”, n = 21) were freshly prepared from blood collected on March 25^th^. A panel of biobanked sera was also obtained from a second cohort (named “pets pre-epidemic” n = 58), composed of animals sampled before the COVID-19 pandemic - between October 2015 and October 2018, containing 51 dogs from 32 popular breeds and seven cats including five European Shorthair, one Turkish Angora and one Devon rex.

The search for antibodies against SARS-CoV-2 was carried out on the 79 sera of the two cohorts using a Luciferase Immuno-Precipitation System (LIPS) assay as described (1), using two antigens: (*i*) the S1 domain of the SARS-CoV-2 Spike S protein and (ii) the C-terminal part (residues 233-419) of the SARS-CoV-2 N nucleoprotein. Viral antigens were produced in HEK-293F cells transfected with plasmids expressing the gene for Nanoluc fused to the C-terminus of the viral protein. Recombinant proteins were harvested from the supernatant (S1, which is a surface glycoprotein secreted in culture supernatant) or cell lysate (N, which is located inside the virus) without any purification step, and incubated with 10 µL of animal serum. The immune complexes were precipitated with protein A/G-coated beads, washed, and the luciferase activity was monitored on a Centro XS^3^ LB 960 luminometer (Berthold Technologies, France). The positivity threshold was defined as the mean of 10 negative controls (without serum) + five standard deviations. Human patients’ sera collected before and during the course of the epidemic were used as negative and positive controls respectively. SARS-CoV-2 specific antibodies were not detected in the “pets epidemic” cohort, and no statistical difference was observed compared to the “pets pre-epidemic” cohort (Figure 1). In addition, nasal and rectal swabs were taken during one week, starting from the day of blood sampling (25 March) and all animals tested negative for the presence of SARS-CoV-2 by RT-PCR (who.int/docs/default-source/coronaviruse/whoinhouseassays.pdf). We concluded that none of the animals included in this study had been or was infected by SARS-CoV-2, despite repeated close intra-species daily contacts on the campus, and more importantly despite frequent and lasting contacts with COVID-19 patients confined to small rooms.

**Figure 1:**
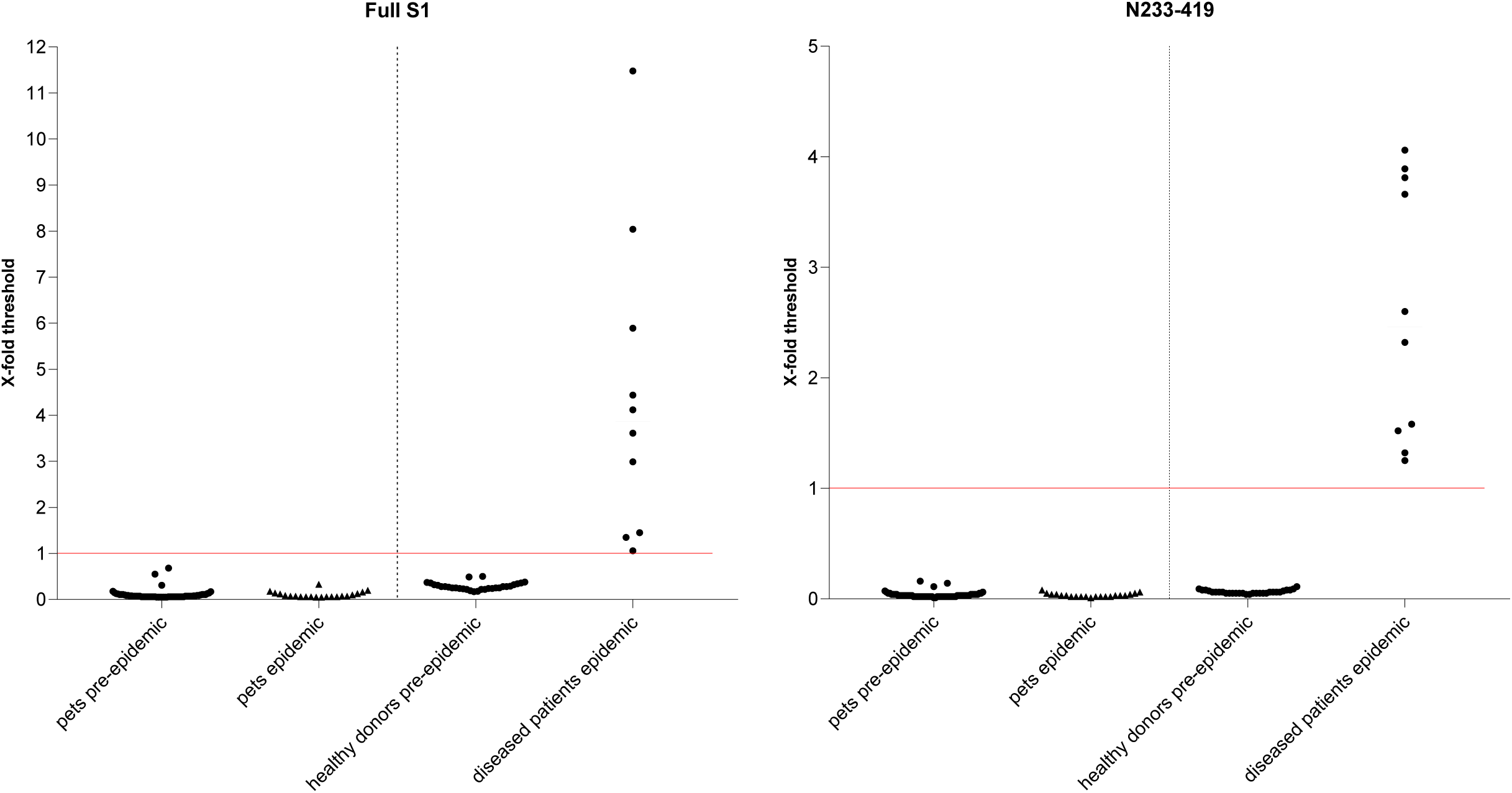
LIPS assay targeting antibodies against SARS-CoV-2 in domestic cats and dogs in close contact with a cluster of COVID-19 patients. Sera from pets in contact with COVID-19 patients (pets epidemic, n = 21); from pets sampled before the epidemic (pets pre-epidemic, n = 79) were tested for the presence of antibodies directed to S1 (left panel) and the C-term (aa 233-419) part of the SARS-CoV-2 nucleoprotein (right panel), using a luciferase immunoprecipitation system assay (LIPS). The same assay was performed for humans unrelated to this study, either COVID-19 human patients (symptomatic human patients epidemic, n = 10) or healthy volunteers blood donors (healthy donors pre-epidemic; n = 30).

A few studies have investigated the putative role of domestic animals in the current COVID-19 pandemic. Of note, Shi *et al*. (3) have demonstrated that among seven domestic animal species tested, the ferret, cat and dog could be experimentally infected by the intranasal route, probably through the viral receptor angiotensin-converting enzyme 2 (ACE2). Contrarily to dogs and ferrets, which were less affected, cats have been shown to be more susceptible to experimental infection, particularly during their juvenile post-weaning life (70-100 days). In these experimental conditions, Shi *et al*. also reported that one out of three naïve cats exposed to infected cats became infected (3). This revealed that intra-species respiratory droplet transmission can occur in cats, at least from cats experimentally infected with 10^5^ pfu by the intranasal route (3). Consistent with the increased susceptibility of this domesticated species, a recent study of 102 pet cats living in Wuhan, China, and collected in the first quarter of 2020 during the pandemic period, reported a prevalence of 14.7% seroconverted cats (4), presumably adult cats.

Although based on a limited number of animals, in particular cats, our study points to undetectable interspecific transmission of the SARS-CoV-2 virus between COVID-19 patients and domestic dogs or cats under the natural exposure conditions. Given the susceptibility of cats inoculated intranasally with 10^5^ pfu or by direct aerosol transmission from infected cats (3), it is conceivable that the infected and seroconverted cats identified in Wuhan, China (4), had either been in contact with patients whose viral load was higher than in our study, or have had more contacts with infected cats. These are questions among others that will need to be further addressed.

Therefore, despite the susceptibility of some animal species revealed in experimental conditions and making the juvenile cat, in particular, a potential model of SARS-CoV-2 replication, our results support evidence for a nil or very low COVID-19 rate of infection in companion dogs and cats, even in a situation of repeated contacts and close proximity to infected humans. This suggests that the rate of SARS-CoV-2 transmission between humans and pets in natural conditions is probably extremely low, below a reproduction number of 1. Replication studies to accurately characterize a possible role of pets, and especially feline, as an intermediate host vector of SARS-CoV-2 are needed in different international contexts. They should include larger populations and the study of the role of the age of the animals and the surrounding viral load.

## Acknowledgements

we thank the pets’ owners who gave us the authorization of sampling, Gwenaelle Leseigneur (veterinary nurse at the EnvA hospital for small animals) for her excellent technical help. We also thank funding from Institut Pasteur, IBEID Labex, Reacting, Cani-DNA BRC that is part of the CRB-Anim infra-structure, ANR-11-INBS-0003 and UE RECOVER

